# TF-IDF k-mer–based Classical and Hybrid Machine Learning Models for SARS-CoV-2 Variant Classification under Imbalanced Genomic Data

**DOI:** 10.64898/2026.04.02.716024

**Authors:** Nazmul Haque, Abdul Mazed, Jannatul Nayeem Ankhi, Md Jamal Uddin

**Affiliations:** Department of Statistics, Shahjalal University of Science and Technology, Sylhet-3114, Bangladesh; Faculty of Graduate Studies, Daffodil International University, Savar, Dhaka – 1216, Bangladesh

**Keywords:** Genomic, Rare variant detection, TF-IDF, Machine learning, Bangladesh

## Abstract

Accurate classification of SARS-CoV-2 genomic variants is essential for effective genomic surveillance, yet it is challenged by extreme class imbalance, limited representation of rare variants, and distribution shifts in real-world sequencing data. In this study, we employed hybrid RF-SVM framework designed for robust detection of rare SARS-CoV-2 variants. It integrates a random forest and a polynomial-kernel based support vector machine to enhance sensitivity to minority classes while maintaining overall predictive stability. We systematically compared classical machine learning models, deep learning approaches, and hybrid strategies under both standard and distribution-shifted evaluation settings. Our results show that classical models using TF-IDF–based k-mer features outperform deep learning methods on macro-averaged performance metrics. The Random Forest classifier using TF-IDF Feature achieved the best overall performance, with a macro-averaged F1-score of 0.8894 and an accuracy of 96.3%. The model also demonstrated strong generalization ability, as evidenced by stable cross-validation performance (CV accuracy = 0.9637). Hybrid RF-SVM model further improves rare variant detection under severe class imbalance. Calibration analysis indicates reliable probability estimates for common variants, although challenges persist for minority classes. Overall, this study highlights the limitations of deep learning in highly imbalanced genomic settings and demonstrates that carefully designed hybrid machine learning approaches provide an effective and interpretable solution for rare SARS-CoV-2 variant detection.

## 1. Introduction

The COVID-19 pandemic caused by Severe Acute Respiratory Syndrome Coronavirus 2 (SARS-CoV-2) has become a worldwide crisis that has caused immense pressure on the healthcare systems of multiple countries [1,2]. By the end of 2025, the accumulated effect had hit an unbelievable rate, showing registered infections of over 775 million and worldwide deaths of over 7 million[3,4]. The initial lab-confirmed cases in Bangladesh were recorded on March 8, 2020 whose footprint had hit Approximately 2.05 million confirmed infections and 29,531 deaths[5,6,7]. One of the main contributors to such high mortality rates and long-term international chaos due to an impressive level of genomic plasticity of SARS-CoV-2[8,9]. These ongoing changes in the genome have not only questioned the effectiveness of current clinical interventions but also has forced the rebalancing of the universal health policies on a daily basis to address the current risks.

Genomic surveillance is a significant instrument and has taken the center stage of controlling global epidemics due to the ability to monitor and track the virus spread[8,10]. Surveillance must be effective, and that is only possible with the correct classification of viral lineages, which enable the investigators to trace the transmission patterns and detect the emergence of Variants of Concern (VOCs). In addition to tracking, it is also necessary to model this genomic data to remain ahead of the pandemic. The modeling assists in the initial identification of novel variants which enable health authorities to be ready before a fresh wave begins. Moreover, the knowledge of such genetic modification can be used to enhance early diagnosis and treatment interventions[11]. When we are aware of the variant in circulation, we are able to select the most effective medicines and this eventually leads to the reduction of risks and saving of lives.

Powerful sequencing Techniques have generated a colossal volume of genomic information that has provided a better view of the genome than ever[12]. Nevertheless, these DNA sequences are hard to analyze since the information is large, complicated, and is usually filled with noises or errors[12,13]. The conventional means are just too slow to handle this amount of information. It is here that machine learning and, particularly, deep learning has transformed the field[12,13,14]. The fact that deep learning is fast and incredibly precise has enabled researchers to classify sequences in a very fast way, which has resulted in an overall improved understanding of how various components of the genome actually operate[12]. A limitation of this real-world genomic dataset is its long-tailed distribution, in which a small number of predominant lineages account for the majority of sequences, while many lineages are severely under-represented [15,16].

Although most consider that deep learning (DL) models are innately better at genomic tasks because they are capable of modeling complicated sequences, the situation is more subtle. Practically, the extreme class imbalance can be a major problem in DL models, which do not learn discriminative features on minority classes, and therefore perform disastrously at macro-average performance[17,18,19]. In addition, genomic pipelines in practice are hardly ever ideal; they are often prone to variation in the quality of sequencing which cuts off or truncates a sequence[19,20]. Such distribution shifts compel conventional models, especially models based on deep representation learning, to trade-off robustness, leading to their accuracy plummeting when they are not tested in the laboratory but in noisy real-world settings[19]. Moreover, LSTMs architecture, as well as other, is notorious in its alleged learning struggle under the condition that training data is small, since they need large datasets to achieve optimal performance. This means that in cases, where certain viral variants are under-represented in the data, the latter tend to overgeneralize and therefore rare lineages effectively disappear[19].

To address these gaps, we use a Hybrid RF-SVM framework that is designed to be more successful when deep learning fails. This framework does not make use of data-intensive neural architectures, rather it uses a dual-classifier approach, which combines TF-IDF k-mer descriptors with the complementary advantages of Random Forests and Polynomial-kernel Support Vector Machines (SVMs). Our method has a high sensitivity of SVMs with a stable probability calibration of Random Forests, making it sharper in detecting rare variants that traditional models can easily overlook. Based on the SARS-CoV-2 genomic data of Bangladesh, we would like to utilize the sparse 6-mer encoding and the dual-classifier architecture to identify the robust genomic features. The aim of this method is to have a strong alternative to surveillance in local clinical conditions, where new variants are frequent and hard to detect because of the lack of data and technical noise.

## 2. Methodology

### 2.1 Study Overview

This study comparing the performance of classical machine learning, deep learning, and hybrid modeling aims to classify genomic variants in extreme class imbalance and distribution shift. By using SARS-CoV-2 entire-genome sequences in Bangladesh, (i) discriminative genomic features were extracted, (ii) a series of predictive models were trained, (iii) robustness against realistic surveillance conditions was measured, and (iv) calibration and ability to detect rare-variants were assessed.

### 2.2 Dataset Description and Quality Control

The data included SARS-CoV-2 genomic sequences in Bangladesh with variant lineage. The filtering of all sequences was done to eliminate ambiguous characters, non-genomic artefacts and only retained canonical nucleotides (A, C, G, T). Variants with fewer than two occurrences in the dataset were considered rare variants. To avoid record leakage and over-predicted performance, all the sequences that were exact duplicates were detected and removed pre-train experiment. Subsequent clean sequences were analyzed in terms of nucleotide composition, GC content and length distribution so as to create a background of compositional conservation. Once cleaned, the data exhibited high class imbalance, with few dominant variants and a long tail of rare lineages, which reflects real-world genomic surveillance data.

### 2.3 Feature Engineering

#### 2.3.1 k-mer TF-IDF Representation

A k -mer-based Term Frequency -Inverse Document Frequency (TF -IDF) encoding was used to encode genomic sequences into numerical representations. Each genome was divided into overlapping k -mers and TF -IDF weighting enhanced to emphasize discriminating patterns in the sequences and mitigated ubiquitous motifs. TF-IDF serves to measure the significance of every k-mer in a document compared to the entire corpus and it is calculated as follows:

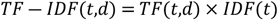

Where,

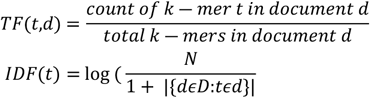

Here, N is the total number of documents, and the denominator counts documents containing the k-mer t.

This representation allows for Efficient encoding of mutation-level information, Robust learning under high dimensionality and Interpretability compared to deep embeddings.

#### 2.3.2 Hand-crafted Genomic Features

Compositional characteristics of the sequences were taken manually and these included the frequency statistics of the nucleotides, the GC content, the average nucleotide composition (percentages of A, C, G and T), and length descriptors of the sequence. These characteristics produce biologically intuitive summarization of genome structure, but do not directly describe positional or contextual mutation patterns.

#### 2.3.3 Merged Feature Sets

A hybrid space of feature was built, which combined TF-IDF features with hand-crafted features that allowed the analysis of the effects of redundancy at the high dimensionality on the model performance.

### 2.4 Data Splitting Strategy

#### 2.4.1 Stratified splitting

The dataset was split into training and test sets using stratified sampling to preserve class proportions. Random seeds were fixed to ensure reproducibility. Additionally, 10-fold cross-validation was performed on the training data for stability assessment.

#### 2.4.2 Hard Split for Distribution Shift

In this conditions, the long sequences only were in trained set and all short sequences and a 15 per cent subset of the long sequences were in tested set, simulating a distribution shift scenario to evaluate its generalization to sequences that differ in length or other characteristics. This simulates real surveillance conditions where sequencing quality varies over time.

### 2.5 Model Architectures

#### 2.5.1 Classical Machine Learning Baselines

##### Random Forest (RF) model

The RF classifiers were trained based on a collection of bootstrap aggregation decision trees. RF was chosen due to its robustness to high dimensional sparse features, ability to capture nonlinear interactions as well as being resistant to overfitting in case of class imbalance. RF models were trained using all combinations of features.

##### Support vector machine (SVM) model

Support Vector Machines were trained using linear kernel, Radial Basis Function (RBF) kernel and Polynomial kernel. The polynomial kernel can demonstrate strong sensitivity to minority classes, particularly under imbalance. SVMs were trained on each feature configuration to evaluate kernel-specific behavior. SVM decision function is as follows:

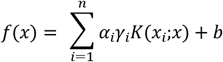

Where, *K*(·) indicates the kernel function.

#### 2.5.2 Deep Learning Models

##### Convolutional Neural Network (CNN)

CNN models were developed to learn hierarchical sequence patterns directly from k-mer–encoded inputs. The architecture consisted of convolutional layers in which the motif is extracted, pooling layers that reduce the dimensionality, and fully connected layers which classify. The CNNs were trained using the backpropagation and early stopping.

##### Long Short-Term Memory (LSTM)

LSTM networks were implemented to model long-range dependencies in genomic sequences. Training was done on sequences that were padded or truncated to constant lengths. Although they can perform sequential modeling, LSTMs were highly scrutinized on small data and imbalance.

#### 2.5.3 Hybrid Modeling Strategies

##### Hybrid Convolutional Neural Network–Random Forest (CNN–RF)

The hybrid CNN-RF was initially trained to acquire hierarchical and localized representations of the sequence using k-mer-encoded genomic input using the Convolutional Neural Network (CNN). Instead of using the CNN as a single-purpose classifier, intermediate representations of the final dense layer were obtained and considered as learned feature vectors. These CNN based features were then inputs to a Random Forest (RF) classifier which did the final classification. This combination approach has the advantage of using the CNN ability to learn informative sequence-level patterns automatically with the strength of the Random Forest to high-dimensional features, noise, and extreme class imbalance. The CNN-RF model is more likely to achieve better generalization than single deep learning models, especially in imbalanced genomic samples; it decouples feature learning and classification to enhance generalization.

CNN feature extraction:

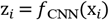

where x_*i*_denotes the k-mer–encoded input sequence and z_*i*_represents the learned CNN feature vector. Random Forest classification:

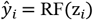

##### Hybrid Support Vector Machine–Random Forest (SVM–RF)

To improve the sensitivity of detection of rare genomic variants a hybrid Support Vector Machine-Random Forest (SVM-RF) model was built by exploiting complementary advantages of both margin-based soft-margin classifiers and ensemble learning paradigms. The former consisted of a Support Vector Machine with a polynomial kernel, which was chosen due to its ability to maximize the margins boundaries in high-dimension feature spaces, which enhances the distinction of the minority classes. At the same time, a Random Forest estimator was fitted to provide strong predictions and high-quality generalization in most variants of the majority. The combination of results between the two classifiers also delivered a balanced trade-off between precision and recall on common and rare classes, thus improving macro-averaged performance metrics and improving the rate at which the rare variants are detected in highly skewed genomic data sets.

Polynomial-kernel SVM decision function:

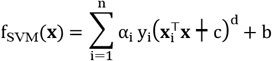

where d is the polynomial degree and c is the kernel offset.

Hybrid prediction integration:

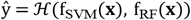

where ℋ (·) denotes the hybrid integration strategy used to balance minority sensitivity and overall stability.

### 2.6 Model Training and Optimization

All models were trained using consistent data splits and random seeds. Hyperparameters were tuned empirically using cross-validation. Early stopping was applied to deep learning models to mitigate overfitting. Convergence behavior, bias-variance trade-offs and sample efficiency are analyzed by generating learning curves.

### 2.7 Evaluation Metrics

Having such an imbalanced classes, the evaluation of model performance was conducted in terms of a variety of measures: accuracy, precision, and recall, F1-score (macro-average and weighted). Especially, the macro-averaged metrics were given priority in order to indicate more on the performance on the minority classes. Class-specific measures were described to ease the rare-variant analysis.

### 2.8 Calibration Analysis

Calibration metrics were calculated to assess the reliability of the probabilities used, and they comprised of the Brier score, Expected Calibration Error (ECE) and Maximum Calibration Error (MCE).

#### Brier Score

The Brier score measures the mean squared difference between predicted probabilities and true class labels:

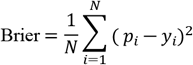

where:

*N* is the number of samples, *p*_*i*_ ∈ [0,1]is the predicted probability for the true class, *y*_*i*_ ∈ {0,1}is the ground-truth label. Lower Brier scores indicate better probability calibration.

#### Expected Calibration Error (ECE)

ECE quantifies the discrepancy between predicted confidence and empirical accuracy by partitioning predictions into *M* confidence bins:

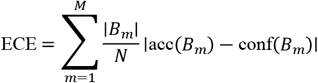

where:

*B*_*m*_ is the set of samples in bin *m*, acc (*B*_*m*_) is the accuracy in bin *m*, conf (*B*_*m*_) is the average predicted confidence in bin *m, N* is the total number of samples. Lower ECE values indicate better calibration.

#### Maximum Calibration Error (MCE)

MCE measures the worst-case calibration error across all bins:

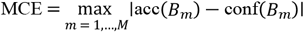

MCE highlights the largest deviation between predicted confidence and observed accuracy.

### 2.9 Reproducibility

All experiments were implemented in Python using Jupyter Notebook environments. The full pipeline including feature extraction, model training, evaluation, and visualization was executed with fixed random seeds to ensure reproducibility. The complete methodological workflow is illustrated in Figure 1.

**Figure 1:**
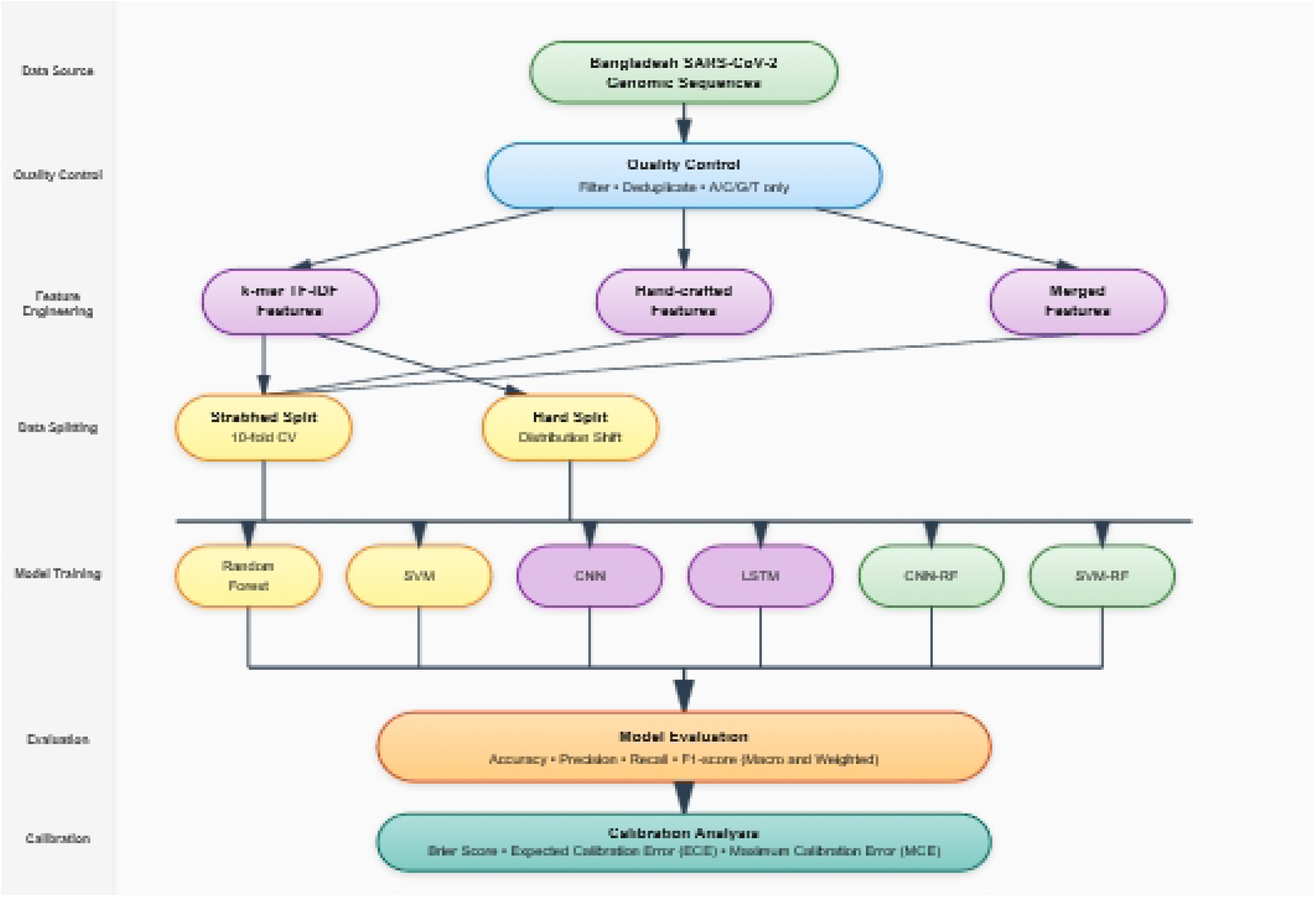
Flowchart of methodological workflow

## 3 Result

### 3.1 Descriptive Analysis

Variant frequency distribution (Figure 2A) showed an extremely imbalanced label distribution. The highest proportions were occupied by lineages 21J (Delta, n = 483) and 20B (n = 451), followed by 21A (n = 288), while many other variants were represented by fewer than 50 sequences. This is typical for the long-tailed character of viral genomic surveillance data. Average nucleotide composition (Figure 2B) was also biased in the direction of adenine (A ≈ 29.8%) and thymine (T ≈ 32.3%) yet contained lower percentages of cytosine (C ≈ 18.3%) and guanine (G ≈ 19.6%). Small standard deviations indicate uniform base composition between genomes. GC content frequency distribution (Figure. 2C) peaked at ∼0.379 (mean = 0.379, SD = 0.006), with interquartile range of 0.373 to 0.380. The tight peak and density overlay suggest strong compositional conservation among sequences with very few outliers below 0.36 or above 0.46. Distribution of sequence length (Figure 2D) revealed most genomes near the predicted ∼29.8 kb (median = 29,803 bp; P95 = 29,903 bp). Some short, truncated sequences (∼3.7 kb at P5) were also observed, corresponding to incomplete or poorly assembled sequences. The inset zoom illustrated the close clustering around the modal length.

**Figure 2:**
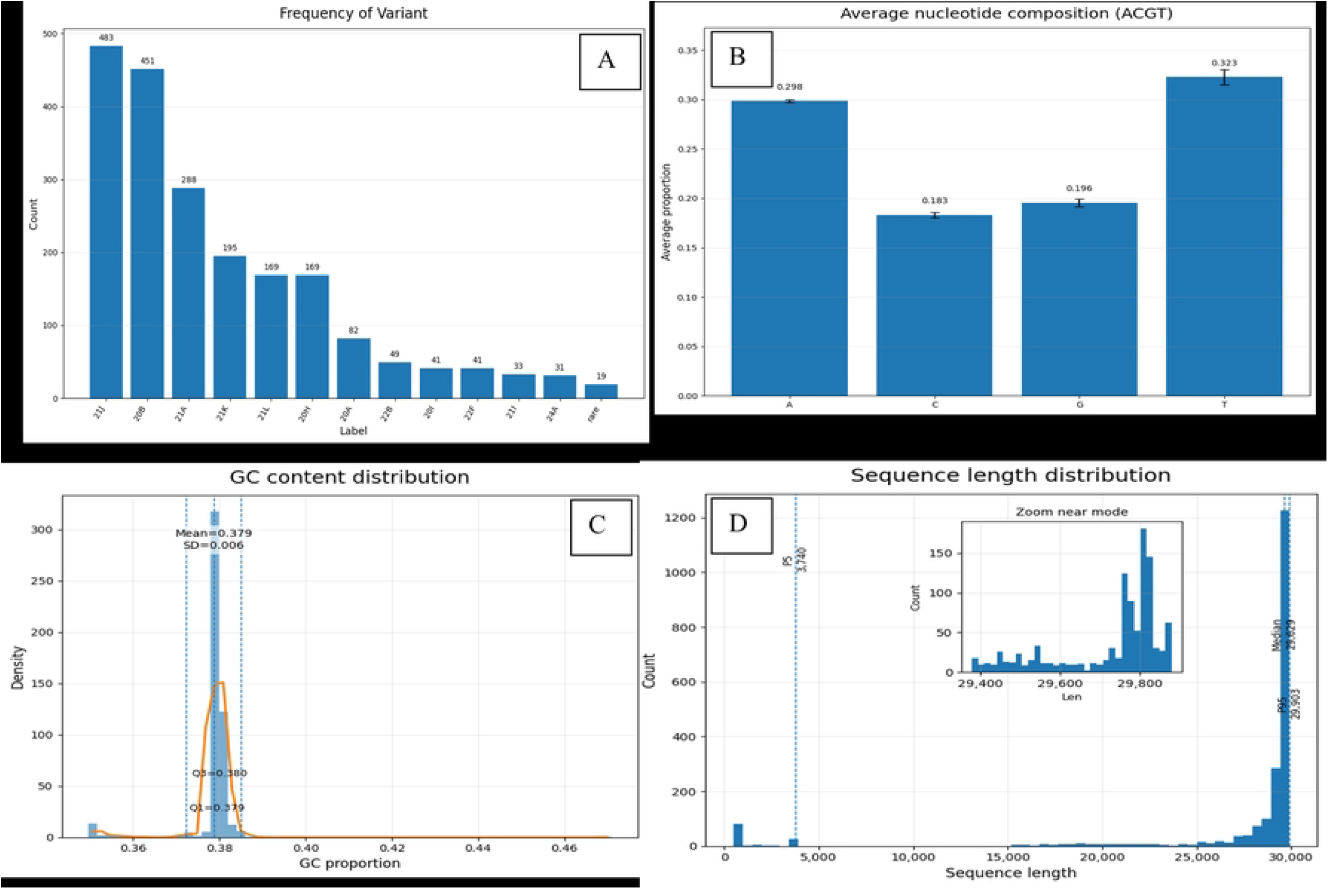
Exploratory analysis ofSARS-CoV-2 genome dataset after cleaning highlighted several sequence properties of special significance.

### 3.2 Baseline Models Performance

We first evaluated baseline models using TF-IDF features. The highest overall performance among them was achieved by Random Forest with a macro-averaged F1-score of 0.8894, 96.3% accuracy, and stable cross-validation performance (CV = 0.9637) in table 1. A similar performance was achieved by the SVM with a polynomial kernel, with a slightly higher macro-average F1-score (0.9007) and good cross-validation accuracy (0.9675), but its overall accuracy (93.3%) was slightly lower than Random Forest. The linear SVM performed middling (macro F1 = 0.7392, accuracy = 89.8%), but the RBF SVM performed poorly, with accuracy plummeting to 25.1% and macro F1 score of 0.3228.

**Table 1:**
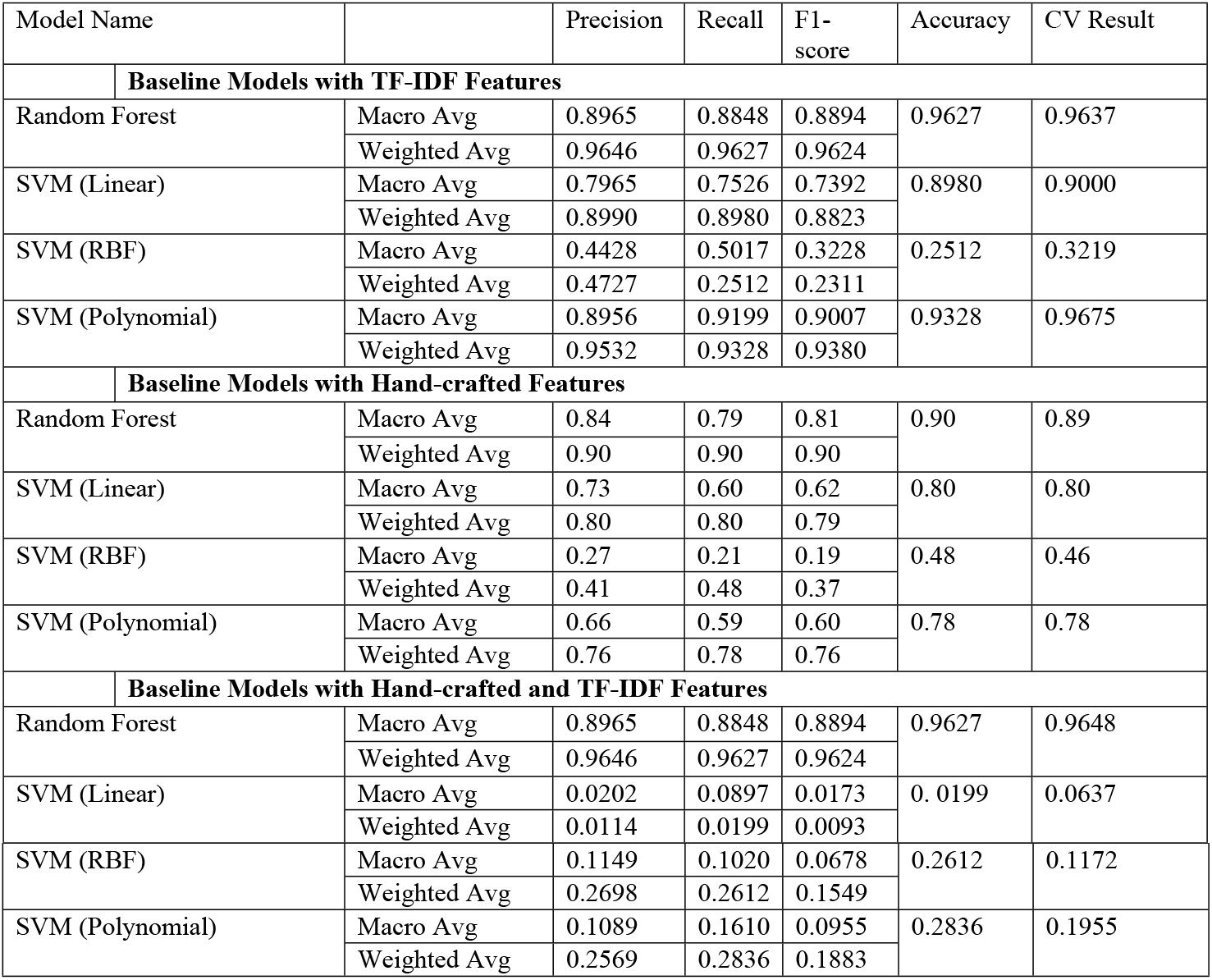
Model performance of baseline models using different features.

Replacing TF-IDF with hand-crafted features, model performance fell across the board. Random Forest operated at a macro F1 of 0.81 and 90% accuracy, while the polynomial SVM achieved a macro F1 of 0.60 and 78% accuracy. The linear and RBF SVMs performed worse on macro F1 (0.62 and 0.19, respectively).

Finally, with all the features merged with TF-IDF again and Random Forest was favored, with continued high performance (macro F1 = 0.8894, accuracy = 96.3%, CV = 0.9648). In contrast, SVM models degraded with this set of features. The linear SVM collapsed (macro F1 = 0.0173, accuracy = 2%), and RBF and polynomial kernels also did worse (macro F1 = 0.0678 and 0.0955, respectively).

### 3.3 Deep Learning Model

From the table 2 it is reported that CNN performed a total accuracy of 71.3%, with a macro-average F1-score of 0.42. While weighted averages had comparatively better performance (F1 = 0.66), the macro-average reveals that the CNN could not classify minority classes well.

**Table 2:** Model performance of deep learning and hybrid (CNN-RF) models.

| Name |  | Precision | Recall | F1-score | Accuracy |
| --- | --- | --- | --- | --- | --- |
| CNN | Macro Avg | 0.40 | 0.45 | 0.42 | 0.7129 |
|  | Weighted Avg | 0.62 | 0.71 | 0.66 |  |
| LSTM | Macro Avg | 0.1276 | 0.1720 | 0.1169 | 0.3060 |
|  | Weighted Avg | 0.2847 | 0.3060 | 0.2213 |  |
| CNN + RF | Macro Avg | 0.8898 | 0.8550 | 0.8681 | 0.9625 |
|  | Weighted Avg | 0.9576 | 0.9625 | 0.9589 |  |

The LSTM performed considerably worse, with only 30.6% accuracy and a macro F1-score of 0.117, which reflects a drastic learning struggle for discriminative features given the data size and imbalance.

In contrast, the hybrid CNN + Random Forest model greatly improved performance by combining the sequence feature extraction power of CNN and the robust classification power of Random Forest. The hybrid model achieved a macro F1-score of 0.8681 and accuracy of 96.3%, which was extremely close to the best achieved using TF-IDF–based Random Forest while still retaining the ability to model sequential k-mer features. Weighted averages were also extremely high (F1 = 0.9589), indicating robust performance across classes.

### 3.4 Rare Variant Detection using Hybrid CNN + RF

Table 3 shows that the Random Forest model performed well overall, except for classes 20H, 21A, 22F, and the rare class, where the SVM model achieved better results. It is particularly notable that for rare variant detection, Random Forest performed very poorly, whereas the SVM model was able to capture nearly 50% of the cases, with an F1-score of 0.500. None of the other models were able to classify the rare variants (see Supplementary File). To address this, we developed a hybrid SVM+RF model, which achieved slightly higher overall accuracy of around 97%, outperforming all other models reported in Table 4. Most importantly, this hybrid model was able to capture the rare variant pattern, as reflected in its precision of 0.500, recall of 0.250, and F1-score of 0.333 for the rare class (support = 4) in table 5. also improved the performance for classes 20H, 21A, and 22F, achieving F1-scores greater than 0.93 for all three.

**Table 3:** Performance metrics (Class-wise) for Random Forest model and Support Vector Machine with polynomial Kernal.

|  | For Random Forest Model |  |  | For SVM (Polynomial Kernel) |  |  |  |
| --- | --- | --- | --- | --- | --- | --- | --- |
|  | precision | recall | f1-score | precision | recall | f1-score | support |
| 20A | 0.8667 | 0.9286 | 0.8966 | 0.4286 | 0.8571 | 0.5714 | 14 |
| 20B | 1.0000 | 0.9888 | 0.9944 | 0.9730 | 0.8090 | 0.8834 | 89 |
| 20H | 0.9375 | 0.8333 | 0.8824 | 1.0000 | 0.8889 | 0.9412 | 18 |
| 20I | 1.0000 | 1.0000 | 1.0000 | 0.8571 | 1.0000 | 0.9231 | 12 |
| 21A | 1.0000 | 0.8846 | 0.9388 | 1.0000 | 0.9038 | 0.9495 | 52 |
| 21I | 1.0000 | 1.0000 | 1.0000 | 1.0000 | 1.0000 | 1.0000 | 5 |
| 21J | 0.9217 | 1.0000 | 0.9593 | 0.9550 | 1.0000 | 0.9770 | 106 |
| 21K | 1.0000 | 1.0000 | 1.0000 | 1.0000 | 1.0000 | 1.0000 | 35 |
| 21L | 1.0000 | 1.0000 | 1.0000 | 1.0000 | 1.0000 | 1.0000 | 36 |
| 22B | 1.0000 | 1.0000 | 1.0000 | 1.0000 | 1.0000 | 1.0000 | 5 |
| 22F | 1.0000 | 0.8667 | 0.9286 | 1.0000 | 1.0000 | 1.0000 | 15 |
| 24A | 0.9286 | 1.0000 | 0.9630 | 0.9286 | 1.0000 | 0.9630 | 13 |
| rare | 0.0000 | 0.0000 | 0.0000 | 0.5000 | 0.5000 | 0.5000 | 2 |

**Table 4:** Performance metrics of Hybrid SVM-RF model.

| Name |  | Precision | Recall | F1-score | Accuracy |
| --- | --- | --- | --- | --- | --- |
| **SVM+RF** | Macro Avg | 0.9314 | 0.9115 | 0.9168 | 0.9708 |
|  | Weighted Avg | 0.9698 | 0.9708 | 0.9693 |  |

**Table 5:**
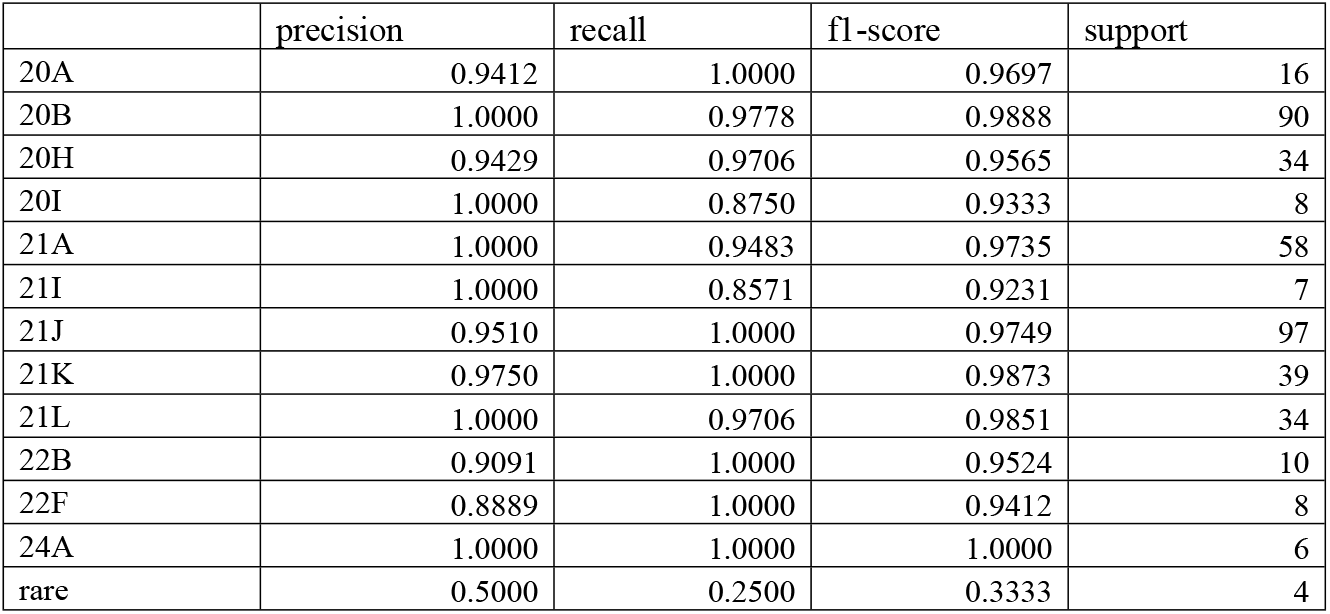
Performance metrics (Class-wise) of Hybrid SVM-RF model.

### 3.5 Hard-Split (Distribution Shift) Evaluation (For Robustness)

In this setting, the training set contained only long sequences, while the test set included all short sequences together with 15% of the long sequences, we can assess that SVM with polynomial kernel achieved the best balance of performance with accuracy of 0.870, recall of 0.834, F1-score of 0.833, and accuracy of 87.5% in table 6. This indicates SVM (Poly) to be much more resilient to distribution shifts than other models.

**Table 6:** Model performance for all models for distribution shift.

| Name | Precision | Recall | F1-score | Accuracy |
| --- | --- | --- | --- | --- |
| **RF** | 0.8204 | 0.7168 | 0.7366 | 0.8158 |
| **SVM (Poly)** | 0.8697 | 0.8344 | 0.8334 | 0.8750 |
| **SVM (Linear)** | 0.6675 | 0.6485 | 0.6416 | 0.7961 |
| **CNN** | 0.5494 | 0.5804 | 0.5277 | 0.6250 |
| **LSTM** | 0.3837 | 0.3137 | 0.3201 | 0.4189 |
| **CNN+RF** | 0.8176 | 0.6019 | 0.6573 | 0.7632 |

Random Forest model remained in competition, achieving 81.6% accuracy and an F1-score of 0.737, though its recall dropped more sharply than SVM. The linear SVM gave average performance (accuracy = 79.6%, F1 = 0.642).

But it is noticeable that deep models performed poorly under this test. CNN performed only 62.5% accurate with F1-score 0.528, and the LSTM performed the worst of all (accuracy = 41.9%, F1 = 0.320). The CNN+RF model also performed lower than RF alone, with accuracy of 76.3% and F1-score of 0.657, which means the CNN component sacrificed robustness in shifted data.

### 3.6 Selected model diagnosis

To ensure reliability, we first verified that the training and test datasets contained no duplicate sequences. We then performed 10-fold cross-validation, which produced consistent results, further supporting the robustness of our findings. Learning curves indicated that Random Forest using 6-mer TF-IDF descriptors achieved nearly optimal accuracy (>99%), enabling strong generalization on majority classes in figure 3. Balanced metrics, however, revealed that RF and SVM met at relatively moderate performance (∼0.82– 0.85 macro-F1), while rare-class F2 stayed low (∼0.25), reflecting continued challenges with minority variant detection.

**Figure 3:**
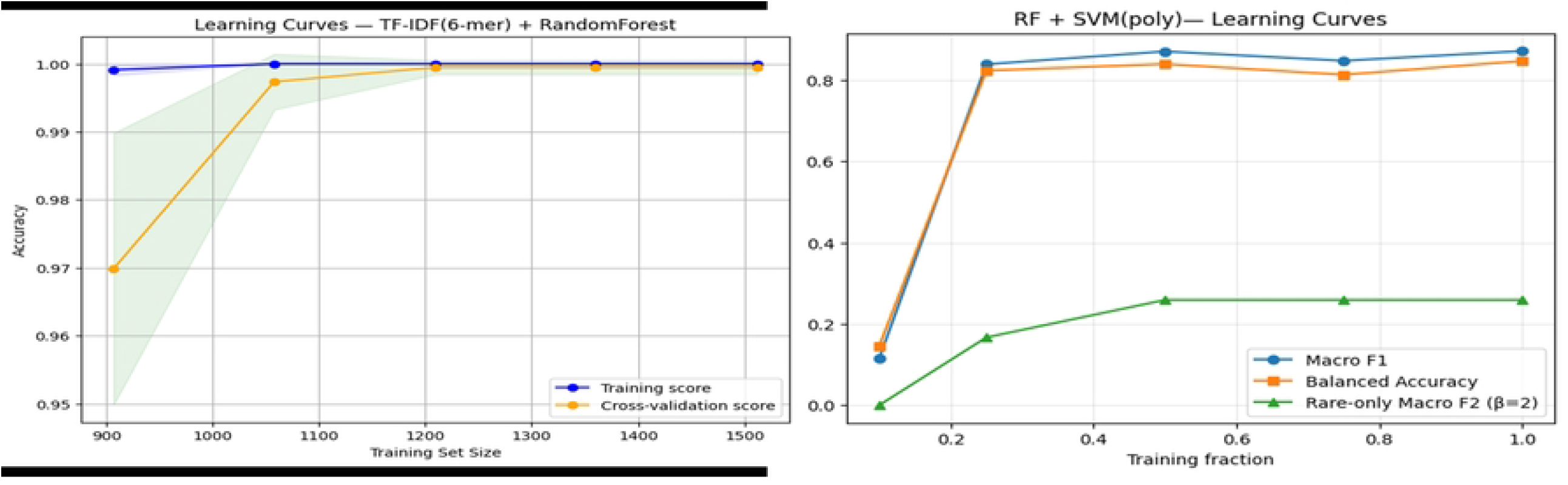
Learning Curves for Random Forest and Hybrid RF-SVM Model.

These results are congruent with the aim of our research in designing a hybrid method: Random Forest offers overall accuracy and robustness, whereas SVM offers higher sensitivity towards minority classes. Together, they pave the way to the achievement of balanced performance in real-world class imbalance and distribution shift.

Calibration analysis showed that the Random Forest and hybrid RF-SVM model shows the identical result based on calibrating matrix in table 7 and figure 4. Both achieved the lowest Brier (0.004) and ECE (0.006) values, indicating well-calibrated probability estimates for common classes. However, the maximum calibration error remained high across all models, reflecting unstable probability reliability for rare variants. The SVM exhibited poorer calibration overall, and the optimal RF+SVM blend defaulted to RF-only, further emphasizing that Random Forest provided the most reliable probability estimates in this setting. The class-wise calibration results are provided in the Supplementary File.

**Table 7:** Macro calibration metrics.

|  | Brier | ECE | MCE |
| --- | --- | --- | --- |
| RF | 0.0040 | 0.0064 | 0.4435 |
| SVM | 0.0188 | 0.0272 | 0.3848 |
| Blend (RF and SVM) | 0.0040 | 0.0064 | 0.4435 |

**Figure 4:**
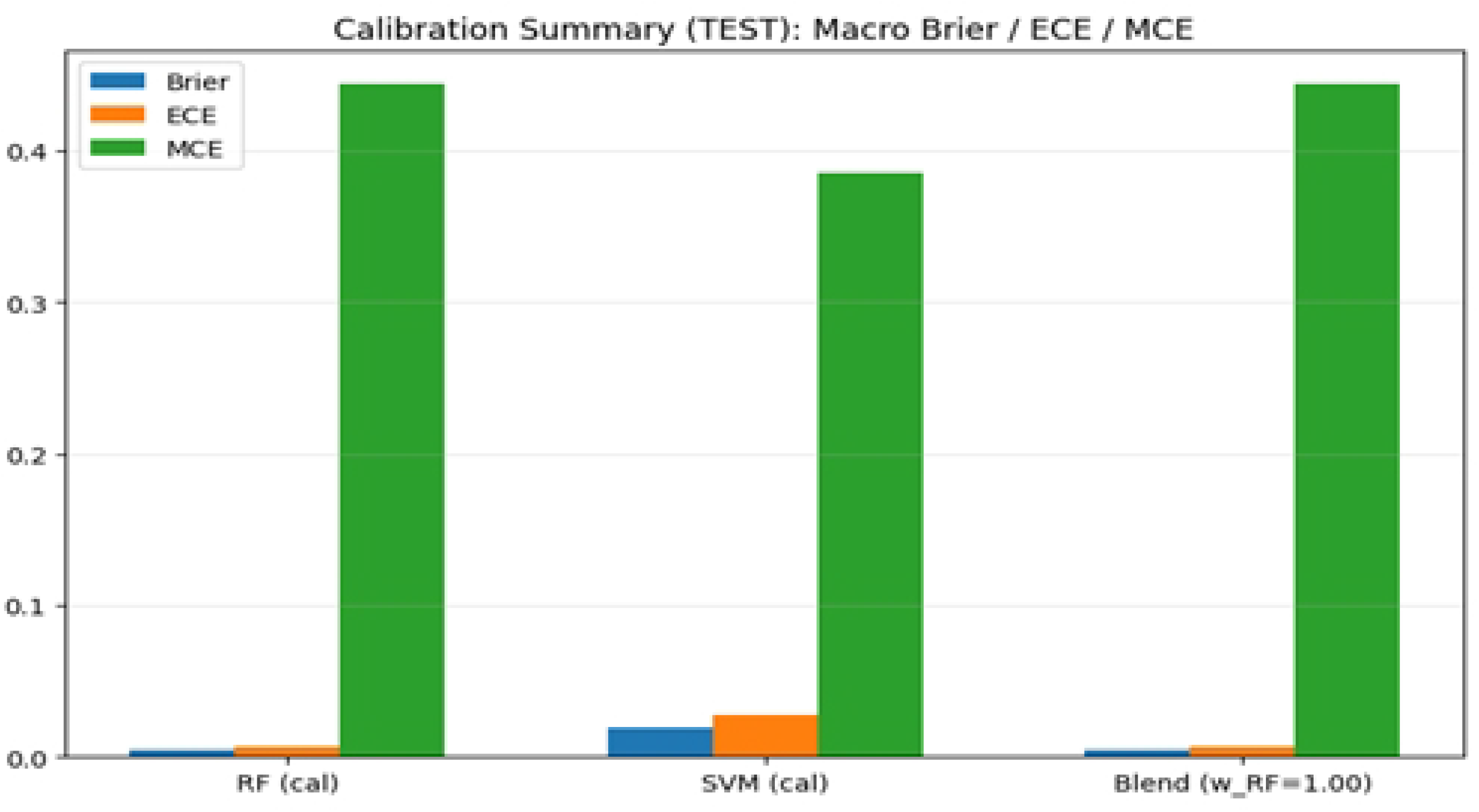
Macro calibration summary.

## 4 Discussion

This paper comparatively studies the classical machine learning, deep learning and the hybrid methods in the context of genomic variant classification with extreme class imbalance. Opposite to the general belief that deep learning models are superior[21,22,23,24,25], the findings reveal that simple baseline algorithms, specifically the Random Forest and the use of the polynomial-kernel support vector machines with TF-IDF k-mer representations, always perform better than the deep neural networks in this case. Such results highlight the significance of the correspondence between model complexity and the nature of data in applied genomics.

An important conclusion of this study is that TF-IDF-based k-mer features are effective. TF-IDF representations had significant better macro-averaged performance than hand-crafted compositional features in all baseline models[26]. This implies that sparse frequency-weighted k-mer data is an effective way to encode discriminative genomic signals even in the face of no explicit sequence-order modelling. Conversely, the addition of a hand-crafted feature to TF-IDF did not enhance the performance and, in certain situations, deteriorated the support-vector-machine performance, which is probably caused by feature redundancy and higher dimension. The findings point to the fact that quality and adequacy of features are more important than quantity and especially in imbalanced and high dimensional genomic data[27].

Classical baselines outperformed deep learning models, such as convolutional neural networks, and long short-term memory networks, particularly when assessed on macro-average metrics. Although weighted metrics indicated the moderate accuracy, macro-averaged F1-scores indicated that deep models failed to classify minority variants. This mismatch is an example of a pitfall of genomic machine learning[28], where weighted averages can hide bad performance on poorly-represented classes. The more specifically poor performance of LSTM models further implies that explicit sequence modelling does not bring benefits in case training data is scarce and class distributions are highly skewed[29].

To some extent, these limitations were alleviated by hybrid modelling[30,31,32]. The CNN-Random Forest model significantly performed better than standalone deep models and also performed as well as the best TF-IDF based Random Forest. This implies that convolutional networks are capable of learning informative low-level sequence features, but the effective use of tree-based classifiers requires one to decide in an imbalanced environment. Nevertheless, the CNN-RF hybrid was found to be less robust to distribution shift which is an indication that CNN derived features could be overfitting to the strong patterns of sequences that dominate the training dataset. Therefore, although hybrid deep models can enhance the accuracy in in-distribution[33], they can result in a loss of generalization in real-life surveillance problems.

Detection of rare variants is still a significant issue in genomic surveillance and model trade-offs between models were important as shown in class-wise analyses. Random Forest performed well with the overall accuracy, but was unable to identify the most rare variants. By comparison, support-vectors-machines based on polynomial-kernel were more sensitive to under-represented classes though with a somewhat lower overall accuracy[34]. When these strengths were combined in the proposed hybrid of SVM and RF, it enhanced the macro-based performance and produced quantifiable improvements in finding rare-variants. Though the F1 scores of rare-class F1 were rather small with very small sample sizes, the corresponding relative improvement in comparison to the baseline models does have a sense in the context of surveillance when it is of utmost importance to identify rare variants at an early stage.

The distribution-shift test also supports the strength of the classical models. Training models on fully assembled genomes and testing them on truncated sequences also led to a decrease in performance in all methods as expected. It is worth noting that the support-vector-machines based on the use of poly-kernel turned out to be the most robust model in the face of distribution shift and performed better than Random Forest, deep learning, and CNN-based hybrids[35]. This indicates that margin based classifiers are more likely to generalize with covariate shift as compared to ensemble or deep representation-learning. The low resilience of deep models in this condition is a worrisome feature of using them in genomic pipelines in real-world applications because the quality and completeness of sequencing often fluctuate.

There were other insights that were realized through calibration analysis. Random Forest and the RF-SVM hybrid obtained the lowest Brier scores and predicted calibration error, which shows that the common variants are well approximated[36]. However, the maximum calibration error was high in all the models which reflects the erratic and overconfident predictions of rare classes. This result points to a significant weakness: the performance of rare-class detectors can be improved without necessarily providing well-calibrated probabilities. This miscalibration has been a major challenge to downstream applications like the assessment of risks to the health of the people.

To conclude, this paper has shown that the simplicity of the model, the right representation of the features, and the strict evaluation are likely to be more crucial than the architectural complexity in imbalanced classification in genome. The findings warn of the blind use of deep learning in environments where few data, high class imbalance, and distribution shift are a challenge. Rather, classical machine learning models, especially when hybrids, provide strong, interpretable, and computationally-efficient alternatives. Further development in the future must be done to enhance calibration of the rare variants, cost-sensitive and Bayesian models, and these hybrid methods can be applied to larger genomic surveillance problems.

## 5 Limitations & Future Work

Although the proposed hybrid RF+SVM method improved sensitivity to rare variants, our calibration analysis found notable limitations in the reliability of predicted probabilities for minority classes. Specifically, while Random Forest produced well-calibrated output for common classes (low Brier and ECE), the maximum calibration error was large due to unstable and overconfident probability estimates in rare classes. This implies that although more rare variants are being detected, the probabilities assigned still cannot yet be fully trusted for downstream risk prediction or clinical interpretation. Closing this gap will require advanced calibration methods, e.g., temperature scaling, Bayesian approaches, or ensemble-based resampling methods, to achieve both accurate rare-event detection and trustworthy probability estimates simultaneously. Future work will also consider the integration of cost-sensitive learning and data augmentation to further alleviate imbalance-induced miscalibration.

## Nextstrain clade (Commonly used name)

20A Early pandemic clade (no WHO name)

20B Early European clade (no WHO name)

20H Beta (B.1.351)

20I Alpha (B.1.1.7)

21A Delta (B.1.617.2)

21I Delta (AY sublineage)

21J Delta (AY sublineage)

21K Omicron (BA.1)

21L Omicron (BA.2)

22B Omicron (BA.5)

22F Omicron (BQ.1 / BA.5-derived)

24A Omicron (XBB lineage)

## Ethics approval and consent to participate

This study was conducted using publicly available, open-access human genomic data related to SARS-CoV-2. All data were obtained from publicly accessible repositories that provide fully anonymized and deidentified datasets. No new human participants were recruited, and no identifiable personal information was accessed or used in this study. Therefore, formal ethics approval and informed consent were not required for this study.

## Consent for publication

Not Applicable

## Acknowledgement

Not Applicable

## Author Contributions Statement

**Nazmul Haque, Abdul Mazed and Jannatul Nayeem Ankhi** jointly conceived and designed the study, prepared visualizations, interpreted the results, and contributed to drafting the full manuscript. **Nazmul Haque** performed statistical analysis and implemented the machine learning models. **Md. Jamal Uddin** Provided guidance on modelling and analysis and contributed to manuscript refinement.

## Data availability statement

All datasets and source code generated during the current study are available in the GitHub repository at https://github.com/NazmulHaque76/HyV-Class-for-SARS-CoV-2-Variant-Classifica

